# ProtFlow: Flow Matching-based Protein Sequence Design with Comprehensive Protein Semantic Distribution Learning and High-quality Generation

**DOI:** 10.64898/2026.02.14.705870

**Authors:** Zitai Kong, Yiheng Zhu, Yinlong Xu, Mingze Yin, Tingjun Hou, Jian Wu, Hongxia Xu, Chang-Yu Hsieh

## Abstract

Designing protein sequences with desired properties is a fundamental task in protein engineering. Recent advances in deep generative models have greatly accelerated this design process. However, most existing models face the issue of distribution centralization and focus on local compositional statistics of natural sequences instead of the global semantic organization of protein space, which confines their generation to specific regions of the distribution. These problems are amplified for functional proteins, whose sequence patterns strongly correlate with semantic representations and exhibit a long-tailed functional distribution, causing existing models to miss semantic regions associated with rare but essential functions. Here, we propose ProtFlow, a generative model designed for comprehensive semantic distribution learning of protein sequences, enabling high-quality sequence generation. ProtFlow employs a rectified flow matching algorithm to efficiently capture the underlying semantic distribution of the protein design manifold, and introduces a reflow technique enabling one-step sequence generation. We construct a semantic integration network to reorganize the representation space of large protein language models, facilitating stable and compact incorporation of global protein semantics. We pretrain ProtFlow on 2.6M peptide sequences and fine-tune it on antimicrobial peptides (AMPs), a representative class of therapeutic proteins exhibiting unevenly distributed activities across pathogen targets. Experiments show that ProtFlow outperforms state-of-the-art methods in generating high-quality peptides, and AMPs with desirable activity profiles across a range of pathogens, particularly against underrepresented bacterial species. These results demonstrate ProtFlow’s effectiveness in capturing the full training distribution and its potential as a general framework for computational protein design.

## 1 Introduction

Proteins are fundamental biomolecules that play crucial roles in organisms, serving as structural components of cells and facilitating various essential biological processes. The design of artificial proteins with desired functionalities has become a cornerstone of modern bioengineering research [18]. However, the vast exploration space of possible protein sequences presents a significant challenge, and traditional biochemical approaches for discovering novel proteins are both time-intensive and expensive. Recently, the deployment of deep generative models has revolutionized protein design by enabling intelligent exploration of the biochemical landscape, offering a more efficient and cost-effective alternative [62].

Given the similarities between protein sequences and natural languages, advanced natural language processing (NLP) techniques, particularly autoregressive (AR) models, represented by GPT [1], have been adapted as a mainstream paradigm for protein sequence design [38]. However, AR models generate sequences in a next-token-prediction pattern, which samples new tokens with higher probabilities based on token sequences already generated. This unidirectional pattern falls short in modeling the long-range dependencies and complex amino acid interactions in proteins, and restricts ARs to represent all possible conditional distributions [27]. Advanced non-autoregressive (NAR) approaches, predominantly diffusion models [15], provide an alternate by simulating the global amino acid interactions in proteins [53]. Despite their advantages, diffusion models partition the transition probability path between the source noise and target data distribution into local and independent segments, resulting in suboptimal paths. Consequently, the generated samples tend to concentrate around high-probability modes of the training data [57, 7, 33]. Upon our in-silico inspections on several contemporary protein sequence generative models, this common issue of distribution centralization also occurs in the case of protein sequences. This issue can significantly influence the modeling of proteins that exhibit long-tailed functional distributions, making it difficult to sample rare or peripheral regions associated with rare but essential functions.

Flow matching (FM) [25] directly learns a continuous and globally consistent optimal probability path that transforms the source noise distribution into the target data distribution, thereby offering a promising solution to the suboptimal path issues in diffusion models [33]. Moreover, FM mitigates the prolonged sampling time of diffusion-based approaches by requiring significantly fewer generation steps. FM has been successfully applied across various domains, including protein structure modeling [60] and nucleic acid sequence prediction [44], yet its potential in protein sequence design remains largely underexplored. A major challenge lies in adapting continuous FM models to accommodate the inherently discrete nature of protein sequences.

Constructing probabilistic paths directly in the high-dimensional discrete symbol space of protein sequences often leads to model degeneracy [5, 25]. Existing approaches often address this issue by introducing discrete relaxation, extending continuous diffusion or FM formulations to discrete settings [5, 3, 44]. However, for functional proteins, whose sequence patterns are strongly correlated with semantic representations, such relaxation may bias modeling toward local compositional statistics of natural sequences, overlooking the underlying global semantic organization. Inspired by the powerful representational capacity of large-scale pretrained protein language models (pLMs) [24, 12], we propose embedding discrete protein sequences into a continuous, biologically meaningful latent space. This embedding provides a smooth protein manifold enriched with biological semantics, thereby enabling FM to fit more structured probabilistic flows and extract deeper biological insights. However, pLM embeddings often suffer from excessively high dimensionality and massive activation issues [48], leading to inefficient use of space and unstable training. To address these challenges, we constructed a semantic integration network to reorganize the representation space of pLMs, facilitating stable and compact incorporation of global protein semantics.

Building on these insights, we present a novel flow matching-based framework **ProtFlow**, designed for comprehensive semantic distribution learning of protein sequences, enabling fast *de novo* sequence generation with high-quality. Operating within the latent space of an advanced large-scale pretrained protein language model ESM-2 [24], ProtFlow employs a rectified flow matching algorithm to efficiently capture the underlying semantic distribution of the protein design manifold. Additionally, we introduced a reflow technique to enable one-step sequence generation. We construct a semantic integration network to reorganize the representation space of ESM-2, facilitating stable and compact incorporation of global protein semantics. We pretrain ProtFlow on 2.6M general peptide sequences and fine-tune it on antimicrobial peptides (AMPs), a widely studied and significant class of therapeutic proteins to combat worldwide antimicrobial resistance problems. AMPs are representative as long-tailed functional proteins since they exhibit unevenly distributed activities across different pathogen targets [54, 35]. Many AMP generative models actually suffer from poor coverage of the training distribution and overfitting to dominant clusters, yielding limited structural novelty and reduced broad-spectrum activity [8]. Experiments show that ProtFlow outperforms various state-of-the-art generative methods in generating high-quality peptides, and realistic, diverse, and novel AMPs with broad spectrum activity, especially against under-represented pathogens in previous studies. These results demonstrate ProtFlow’s effectiveness in capturing the full training distribution and its potential as a powerful new tool to aid in discovering functional proteins that have been unintentionally left out in some of the early proposals of leveraging generative model.

## 2 Methods

In this section, we outline our method illustrated in Figure 1. The motivation of our design is to facilitate comprehensive distribution learning in the protein semantic representation space, which in turn generates high-quality protein candidates with broad functionality coverage. At a high level, ProtFlow (1) converts discrete sequences into a semantically meaningful latent space with a pLM semantic integration network; and (2) learns the optimal probability paths comprehensively modeling the training data distribution with rectified flow matching. The detailed model architecture and hyperparameter settings, training settings, and the compression ratio detection can be found in the **Appendix A-C**, respectively.

**Figure 1.**
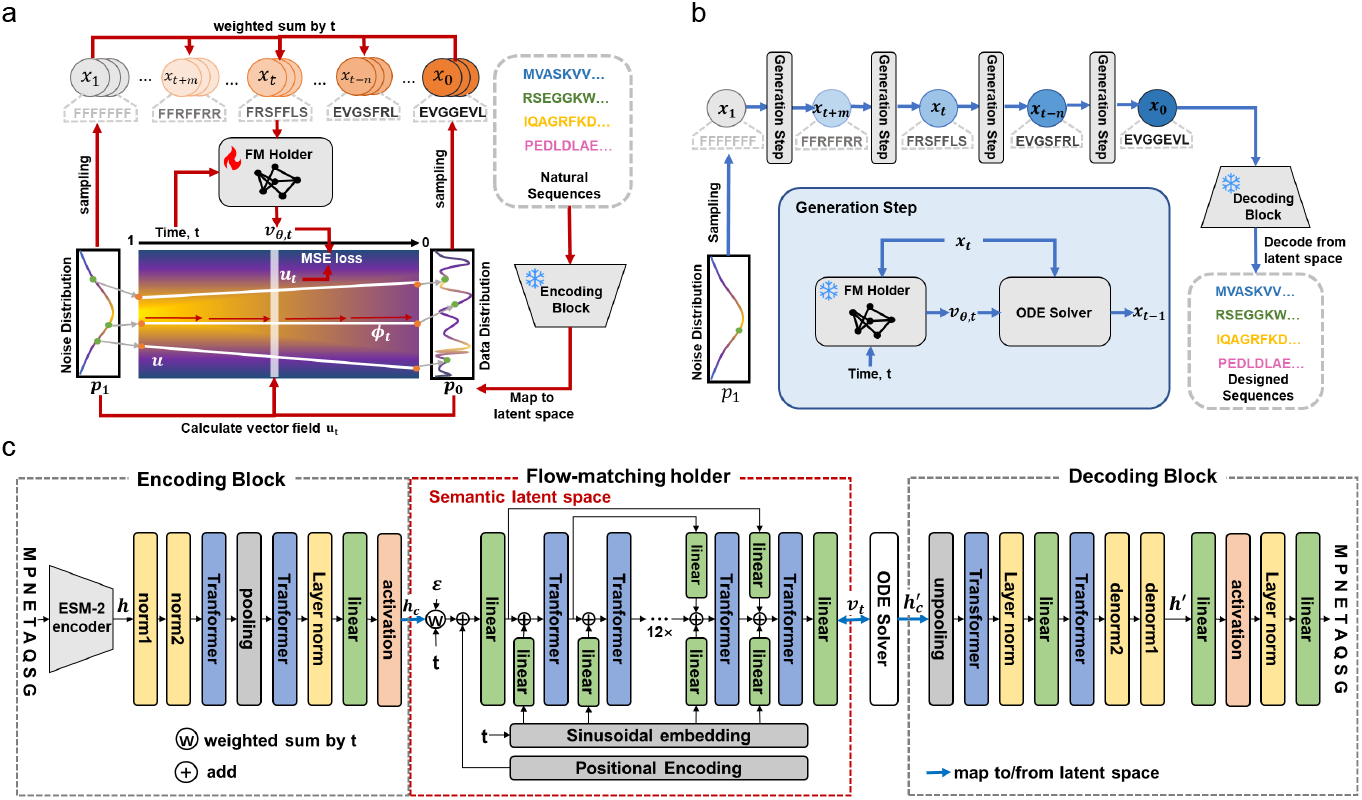
Overview of ProtFlow. (a) Training with the flow-matching algorithm. Standard Gaussian noise and protein embeddings are first sampled. The embeddings and vector fields at intermediate time steps are calculated, and the FM holder learned the vector fields. (b) Inference process of ProtFlow. Initial standard Gaussian noises are first sampled, and the protein embeddings are reconstructed by solving the learned FM vector field through an ODE solver. (c) The model architecture of ProtFlow. ProtFlow involves a pLM semantic integration network consisting of an encoding-decoding block pair, a flow-matching holder and an ODE solver.

### 2.1 Problem formulation

Typically, a protein can be represented as a sequence of amino acids *x* = [*x*_1_, *x*_2_, …, *x*_*L*_] of length *L*, where each amino acid *x*_*i*_ is selected from a vocabulary 𝒱 that includes the 20 standard amino acids. Deep generative models for *de novo* protein sequence design are defined to learn the data distribution *p*(*x*) and to sample novel and plausible protein sequences *x* from this distribution.

### 2.2 Semantically Meaningful Integration to pLM Latent Space

For diffusion and flow matching methods that were originally designed for continuous data types, constructing probabilistic paths directly in the high-dimensional discrete symbol space, such as protein sequences *p*(*x*), often leads to model degeneracy [5, 25]. Considering that proteins are biologically functional molecules whose sequence patterns are strongly correlated with semantic representations, we proposed to map protein sequences *x* = [*x*_1_, *x*_2_, …, *x*_*L*_] ∈ 𝕍^*L*^ into continuous embeddings *h* = [*h*_1_, *h*_2_, …, *h*_*L*_] ∈ ℝ^*L×D*^ in the semantically meaningful latent space of pLMs [17], instead of conventional discrete relaxation [5, 3, 44]. Here, we utilized the encoder of an advanced pre-trained pLM, ESM-2 [24], which is trained on a large-scale standard protein sequence database Uniref50 [45]. We also discussed the effects of different encoders in Appendix D with ablation studies. Through this process, we convert the modeling to a global and underlying semantic distribution context, which may avoid biasing modeling toward local compositional statistics. In addition, we train a corresponding decoder *Dec*_*ψ*_ to reconstruct the sequence as 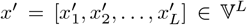. To standardize the length *L* of all protein sequences, we add padding tokens to the ends of shorter proteins, which will implicitly encode the length information in the latent embedding. We utilized the masked cross-entropy reconstruction loss for decoder training.

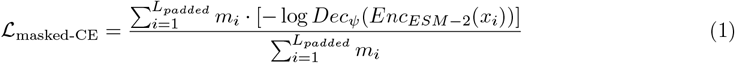

### 2.3 Redesign Latent Space to Enable Compact and Stable Integration

However, pLM embeddings are reported to suffer from problems of unnecessarily large dimensions and massive activation; directly leveraging ESM-2 embeddings may lead to robustness reduction of distribution learning, inefficient use of computing space and training stability [47]. To enhance space efficiency, we introduce a dimensional compression-decompression module pair. This mechanism compresses the original continuous embedding *h* ∈ ℝ^*L×D*^ to *h*_*c*_ ∈ ℝ^*L×D/c*^ with a compression ratio *c* and subsequently reconstructs it back to *h*′ ∈ ℝ^*L×D*^. The discussion of the effects and selection of the compression ratio is in Appendix C. To adjust the data scale and smooth the data, we add preprocessing operation layers before the compression module. Since we use standard Gaussian noises as the source distribution, we apply z-score normalization to scale the output embeddings of ESM-2. The massive activation problem introduces outliers with excessively small variances, which affects the proper convergence. To fix this, we truncate variances with too small absolute values in the z-score using a saturation function and employ a min-max normalization to smooth the truncated latent space.

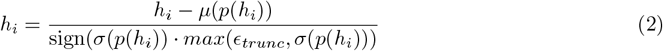

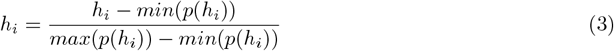

The corresponding reverse operations are added after the decompression block. With the other parts frozen, the compression-decompression pair is trained using a masked MSE reconstruction objective between *h* and *h*′. We also included ablation studies for the decompression architecture in Appendix D.

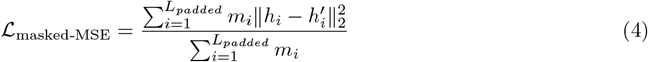

### 2.4 Fast Distribution Learning and Generation with Flow Matching

Inspired by score-based generation models [43], we can define the transition between samples from the data distribution *x*_0_ ∼ *p*_0_ and the prior distribution *x*_1_ ∼ *p*_1_ as an ordinary differential equation

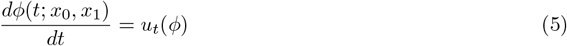

where *ϕ* is an interpolation probability path, namely **flow**, between paired samples (*x*_0_, *x*_1_); *u*_*t*_ is a time-dependent vector field. **Flow Matching** [25] simulates *u*_*t*_(*x*) through a neural network *v*_*θ*_(*x, t*). Then we can train this network by minimizing the loss function:

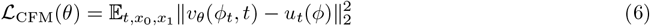

To reduce the complexity of the flow and accelerate the sampling process, **1-Rectified Flows (1-RFs)** [26] define the probability paths as straight lines, where

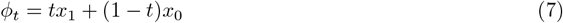

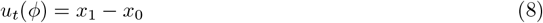

FM provides an effective strategy to overcome the high-probability distribution centralization issue brought by suboptimal path learning or autoregressive generation of other generative methods, while it addresses the long generation time inherent in diffusion-based methods. We expand this Rectified Flow algorithm to the protein sequence design case. Due to its straight and optimal probability paths, our model can theoretically comprehensively modeling the training data distribution and generate good results with fewer ODE-solving steps. During training, we randomly sample the sequence *x*, standard Gaussian noise *ϵ* and time step *t* uniformly within the time range [0, 1]. The sequence will be encoded to *h*_*c*_. Then we can train a flow matching holder network *θ* with the loss function:

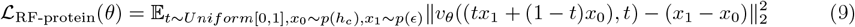

The detailed training procedure is in Algorithm 1 and visualized in Figure 1(a).

During inference, we begin by sampling a random standard Gaussian noise *ϵ* and iteratively solve the ODE within the learned vector field using an ODE solver. The resulting embedding 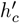 is then decoded back into a protein sequence. For 1-Rectified Flow, we employ the Dormand-Prince45 solver [21] and for the reflow, we use the Euler solver. The optimal number of ODE-solving steps is determined based on specific tasks, with the step range set between [1, 100]. Although the length of protein sequences is implicitly captured in the latent embedding through the trailing padding tokens, we additionally sample the sequence length from the empirical distribution observed in the validation dataset and apply attention masks to ensure the adequate distribution of generated sequence lengths. The detailed inference procedure is in Algorithm 2 and visualized in Figure 1(b).

#### Algorithm 1 Training FM holder with 1-RF

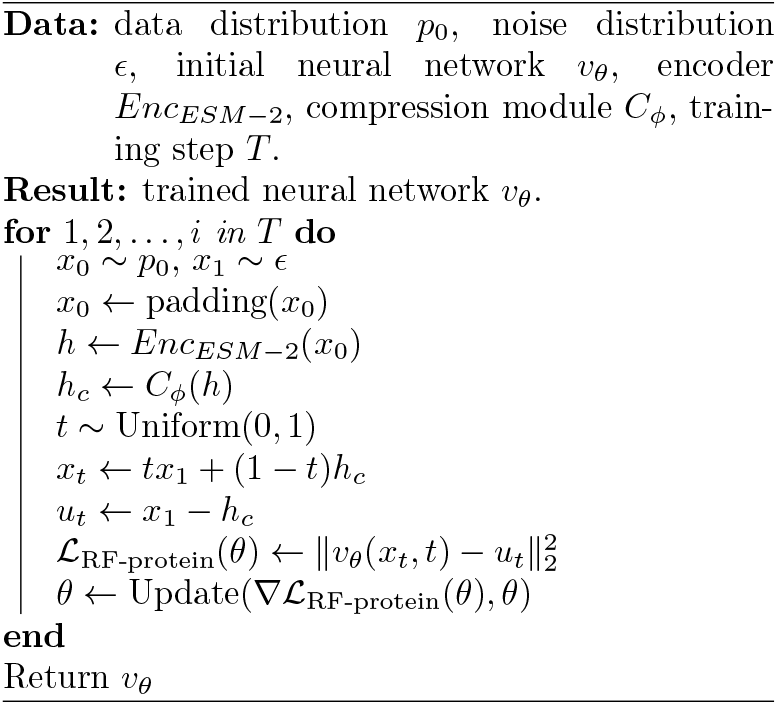

#### Algorithm 2 ProtFlow Sampling

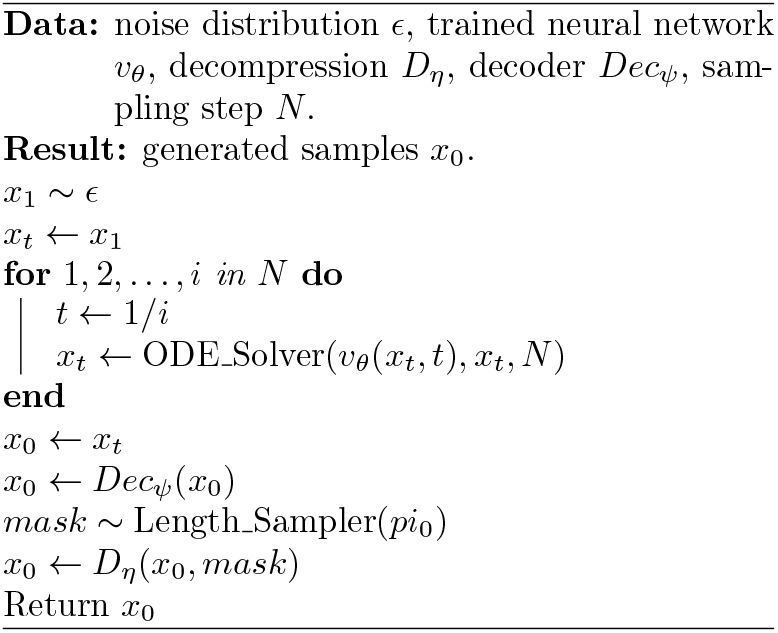

### 2.5 One-step Generation with Reflow

Although rectified flows reduce the transport cost, there are still curves and crossings in the probability paths. To further straighten probability paths, we employ the **Reflow** [26] technique. We sample corresponding pairs (*z*_0_, *z*_1_) with *z*_0_ ∼ *ϕ*_0_(*z*) and *z*_1_ ∼ *ϕ*_1_(*z*) from the start and end point of the ODE solver for the inference of trained 1-Rectified Flow models. The obtained paired noises and data (*z*_0_, *z*_1_) will replace (*ϵ, h*_*c*_) in Equation (9) for finetuning. Rectified Flows without reflow operation are called 1-Rectified Flow (1-RF), and these undergo a single reflow operation are referred to as 2-Rectified Flow (2-RF). With 2-RF, high-quality outcomes can be sampled with only one single ODE solving step. The detailed retraining procedure is in Algorithm 3 and the reference procedure is the same as Algorithm 2.

#### Algorithm 3 Retraining ProtFlow with reflow

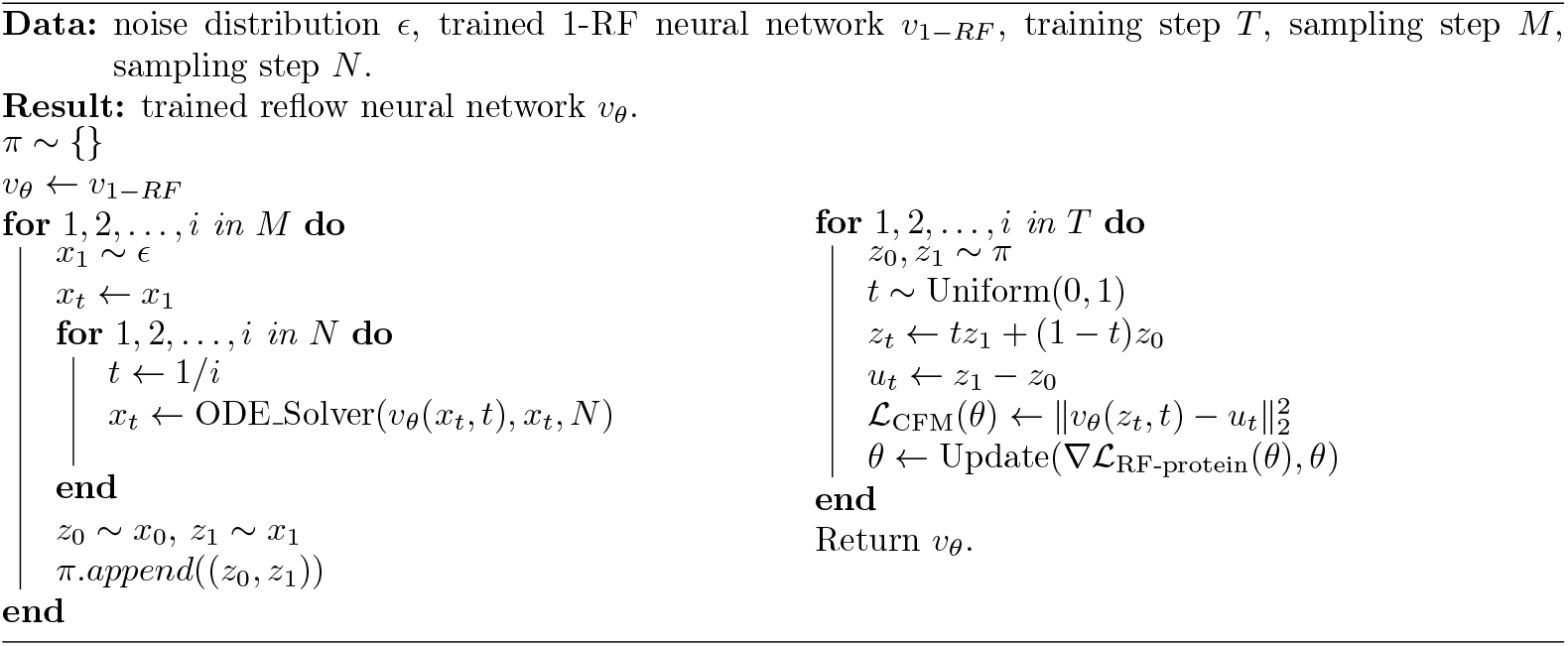

### 2.6 Model Architecture

The diagram of the model architecture is illustrated in Figure 1(c), and detailed hyperparameter settings can be found in the Appendix B. The architecture consists of an encoding block, a flow-matching holder network and a decoding block.

The encoding block consists of an encoder, a dimensional z-score normalization layer with saturation, a min-max normalization layer and a compression module. For the encoder, we use the encoder of ESM-2 35M. For the compression module, we utilize two Transformer layers, a pooling layer, a linear layer and an activation layer.

The decoding block consists of a decompression module, two corresponding denormalization layers, and a decoder. The decompression module has an unpooling layer, two Transformer layers, and a linear layer. For the decoder, we employ the same architecture as the ESM-2 LMHead, which consists of two linear layers, an activation layer, and a layer norm.

The backbone of the flow matching holder module is a BERT model. To match the compressed hidden size, we add two additional linear layers to make projections between hidden size and the compressed hidden size. The noised protein sequence embeddings are added with positional encodings before being fed into the flow matching holder module. Timesteps are projected to match the size of the protein sequence embeddings using a sinusoidal embedding block, and these are added to the input of each Transformer block after a linear projection. Inspired by the U-Vit architecture [6], and following insights from diffusion-based studies [30], we utilize long skip connections, which add the linear projections of earlier block inputs to those of later blocks, to enhance the performance and facilitate more effective learning dynamics.

## 3 Results

Directly training on limited datasets, such as AMPs, leads to low novelty in the generated sequences [52]. Inspired by the idea of fine-tuning models pretrained on large-scale dataset to small dataset [52, 22] of LLMs, we break the workflow of ProtFlow into two stages. ProtFlow was first pre-trained on UniProt peptides to learn the syntax rules and patterns of general peptide sequences, enabling the generation of high-quality non-functional peptides. We then gathered and preprocessed 10k AMP samples from various AMP datasets, and fine-tuned ProtFlow to form the specialized model. For the metrics used, we generally follow two solid works [30] and [52] for thorough evaluations. The supplementary descriptions of ablation study, the details of MIC regressors and top 10 bacterial species selections, baseline selections, definition of metrics can be found in **Appendix D-G**, respectively.

### 3.1 Experimental Settings

#### Datasets

The dataset used in this study comprises three components. For general peptides, we collected proteins with lengths from 2 to 50 from the large-scale standard protein database UniProt [10]. After removing duplicated sequences and sequences with uncommon amino acids (B, U, J, O and X) and lowercase characters, we run easy-search of MMseqs2 clustering with cov-mode 2 and *>* 80% sequence overlap. We split the dataset into 2614953 training peptides and 5270 validation ones following [3].

For AMPs, we collected an AMP dataset from 8 public AMP databases: CAMPR4 [14], APD3 [51], AMP-Scanner v2 [50], GRAMPA [55], DRAMP [40], dbAMP 3.0 [58], DBAASP v3 [34], and StarPep [2]. We keep natural AMP records for CAMPR4 and APD3, general AMPs for DRAMP. and monomer for DBAASP. Similarly, we filtered sequences with lengths between 2 and 50 amino acids, removing duplicates and sequences containing uncommon amino acids and lowercase characters. Subsequently, we performed clustering using Blastp v2.13.0 [59], removing sequences with more than 80% coverage and 60% identity. Finally, we conducted a secondary filtering step using the CD-HIT [23] tool with a threshold of 0.9 to further refine the dataset. This multi-step process ensures a diverse and high-quality dataset for training and evaluation. Ultimately, we obtained 9,938 antimicrobial peptide (AMP) sequences, which were divided into a 8:1:1 ratio for the training, validation and test datasets.

To train prediction models for antimicrobial activity of peptides against various microorganisms, we collected known AMP data with logMIC values from the GRAMPA [55] and DBAASP [34] datasets. We filter these data by selecting sequences with lengths between 2 and 50, removing duplicates and sequences containing uncommon amino acids, and standardizing the expressions of bacterial names. To ensure the training reliability and robustness, we selected the top 10 bacterial species with the most corresponding AMP data. Each specific dataset is split into training, validation and test datasets with a ratio of 8:1:1. **Baselines** For the analysis of general peptide design task, we compared our method with leading methods across different generative model paradigms, including Generative Adversarial Networks (GAN)-based model ProteinGAN [37], AR-based model ProtGPT2 [13], discrete diffusion-based model EvoDiff [3], continuous diffusion-based model DiMA [30], and discrete FM-based model Dirichlet FM [44]. To ensure a fair comparison, all baseline models were configured to have a similar number of parameters as the FM holder of ProtFlow and trained on the same dataset. DiMA used the same ESM-2 encoder as ProtFlow. Each method generated 1,000 peptide sequences for evaluation.

For the analysis of AMP, we incorporated state-of-the-art unconditional *de novo* AMP generative models, including diffusion-based AMP-Diffusion [9], CVAE-based HydrAMP [46], WAE-based PepCVAE [11], and GAN-based AMPGAN [36]. For each model, as well as real AMPs and UniProt, we randomly selected 1,000 protein sequences for analysis.

### 3.2 ProtFlow effectively covers the distribution of natural peptides

We first evaluate the similarity and coverage of the protein spatial distribution to reveal the comprehensive semantic distribution learning ability of ProtFlow. To analyze in a semantically meaningful way, we used another advanced pLM, ProtT5 [12], to convert generated peptides and validation natural sequences into embeddings. To evaluate their distributional similarity from different perspectives, we calculated three metrics: Fréchet ProtT5 Distance (FPD) [3], maximum mean discrepancy (MMD) [41], and 1-Wasserstein optimal transport distance (OT) [42]. FPD is a variant of Fréchet distance (FD) applied to ProtT5 embeddings, which measures the dissimilarity between two samples drawn from multivariate Gaussian distributions. MMD is a kernel-based statistical test to determine whether two sets of samples belong to different distributions. OT leverages the Earth Mover Distance (EMD) solver [31] to determine optimal sequence pairs, and takes the average of the Levenshtein distances [4] between optimal pairs. For all three metrics, lower values indicates greater similarity. ProtFlow consistently achieved significantly lower values across all metrics, demonstrating its ability to generate protein sequences that closely resemble natural proteins in semantic space (Table 1).

**Table 1:**
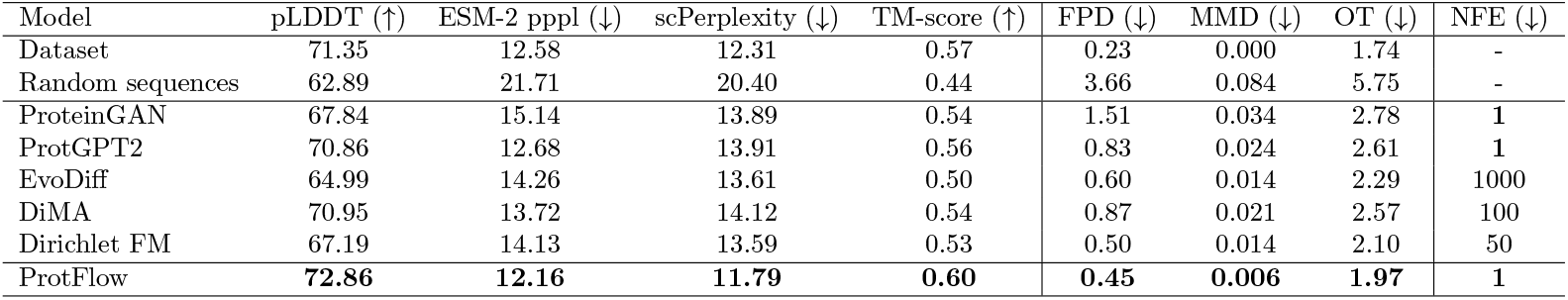
Comparison of computational properties between ProtFlow and baseline models on UniProt general peptides.

| Model | pLDDT (↑) | ESM-2 pppl (↓) | scPerplexity (↓) | TM-score (↑) | FPD (↓) | MMD (↓) | OT (↓) | NFE (↓) |
| --- | --- | --- | --- | --- | --- | --- | --- | --- |
| Dataset | 71.35 | 12.58 | 12.31 | 0.57 | 0.23 | 0.000 | 1.74 | - |
| Random sequences | 62.89 | 21.71 | 20.40 | 0.44 | 3.66 | 0.084 | 5.75 | - |
| ProteinGAN | 67.84 | 15.14 | 13.89 | 0.54 | 1.51 | 0.034 | 2.78 | **1** |
| ProtGPT2 | 70.86 | 12.68 | 13.91 | 0.56 | 0.83 | 0.024 | 2.61 | **1** |
| EvoDiff | 64.99 | 14.26 | 13.61 | 0.50 | 0.60 | 0.014 | 2.29 | 1000 |
| DiMA | 70.95 | 13.72 | 14.12 | 0.54 | 0.87 | 0.021 | 2.57 | 100 |
| Dirichlet FM | 67.19 | 14.13 | 13.59 | 0.53 | 0.50 | 0.014 | 2.10 | 50 |
| ProtFlow | **72.86** | **12.16** | **11.79** | **0.60** | **0.45** | **0.006** | **1.97** | **1** |

To visualize the distribution coverage, we projected embeddings into two dimensions using UMAP [29] and plotted the peptides generated by each model alongside natural peptides(Figure 2d-i). Although all models were trained on the same dataset, we observed that only ProtFlow was able to comprehensively cover the overall distribution of natural peptides, consistent with the distributional relationships revealed by the aforementioned three metrics. It even managed to generate distributions similar to those of the peripheral, protruding, and outlier regions that are farther from the high-density areas. Proteins in these areas can show extreme physicochemical properties, skewed amino acid compositions, and strong structural regularity, making them hard to learn (see Appendix H). In contrast, other models primarily focused on fitting the core regions of high-density distributions.

**Figure 2.**
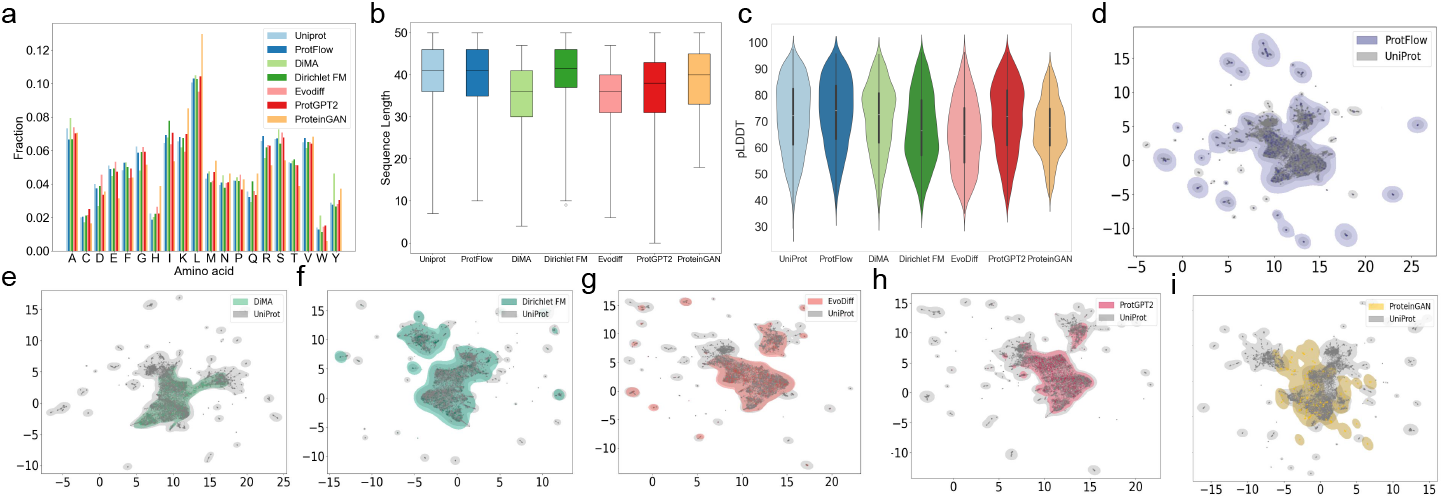
Comparison of physicochemical and distributional properties on designed general peptides. (a) Amino acid frequency distribution; (b) Sequence length distribution; (c)pLDDT distribution; (d)-(i) Natural distributional coverage in protein semantical embedding space.

### 3.3 ProtFlow efficiently generates high-quality peptide sequences

To demonstrate the ability of ProtFlow to generate natural, reliable, and structurally plausible peptide sequences, we perform a multi-level analysis to the generated sequences.

For **sequence-level evaluation**, explicit measurements involve the frequencies of each amino acid and the length distribution (Figure 2a-b). ProtFlow’s results closely matched the natural sequences from UniProt, demonstrating its ability to capture compositional characteristics. However, these metrics primarily reflect superficial sequence properties. To assess semantic-level similarity, we introduced ESM-2 Pseudoperplexity (ESM-2 pppl) (Table 1) with ESM-2 35M by measuring the likelihood of correctly predicting each amino acid in a sequence based on its context. This metric measures how well a given protein sequence aligns with the patterns the assessing model has learned from the large-scale training data of ESM-2. ProtFlow achieves the best perplexity results, revealing that it is capable of generating protein sequences with high reliability and good-alignment with natural protein patterns.

For **structural analysis**, we utilized the mean predicted local distance difference test (pLDDT) score [19] from OmegaFold [56] to assess foldability and structural plausibility (Figure 2c, Table 1). The pLDDT score is a measurement of confidence in the predicted structure. Higher pLDDT scores indicate that the residual interactions within the tested protein are similar to training sequences. Leveraging the structure PDB files predicted by OmegaFold, we use the FoldSeek [20] easy-search tool to identify the most similar proteins in the AlphaFold SwissProt [49] database and calculated the mean TM-Score [61] (Table 1). TM-Score measures global structural alignment, directly reflecting structural similarity. The peptides designed by ProtFlow achieved the highest average pLDDT and TM-Score, demonstrating their structural foldability and structural similarity to natural proteins. At the same time, the TM-score was not excessively high (*<*0.7), indicating that the generated sequences also exhibit good novelty.

To further validate **sequence-structure reliability**, we used ESM-IF 142M [16] to inverse fold the predicted structures back to sequences. We then calculated Self-Consistency Perplexity (scPerplexity) by comparing the inversely folded sequences with the original generated sequences (Table 1). ProtFlow exhibits a leading low scPerplexity, which indicates better consistency between the sequence and its corresponding structure, further validating the reliability of the protein generated from a perspective considering both sequence-level and structure-level.

Notably, flow matching-based methods, ProtFlow and Dirichlet FM, showed advanced performances on distributional metrics over corresponding diffusion models DiMA and EvoDiff, highlighting the fully-covered distributional learning effectiveness of FM-based approaches. Besides, continuous methods lever-aging the latent space of ESM-2, such as ProtFlow and DiMA, consistently outperformed discrete methods like Dirichlet FM and EvoDiff across all metrics. This highlights the advantage of incorporating semantic information from pLMs, which enhances generative model performance by improving the learning of semantic distributions, together with protein sequence and structural characteristics.

### 3.4 AMPFlow generates high-quality AMPs

As functional proteins, the design of AMP requires adequate physicochemical and functional evaluations. We first calculated amino acid frequencies and length distributions(Figure 3a, c). Most naturally discovered AMPs are cationic and amphiphilic, properties critical for inserting into or disrupting bacterial membranes [28]. To further characterize these physicochemical properties, we used modlAMP [32] to visualize total charge, Eisenberg hydrophobicity, and Eisenberg hydrophobic moment (Figure 3b,d,e). To evaluate the similarity in sequence composition between generated and real AMPs, we calculated the 6-mers Jaccard similarity coefficient (JS-6). This metric divides both the training dataset and the generated peptide set into subsequences of length 6, and computes the proportion of overlapping subsequences. Furthermore, to reflect the semantic validity and reliability of generated sequences, we recalculated the ESM-2 pppl (Table 2). ProtFlow exhibit physicochemical property distributions that are highly similar to real AMPs, including similar amino acid frequencies and length distributions, a charge distribution significantly shifted toward higher values, and moderately high hydrophobicity and hydrophobic moment distributions. At the sequence composition and semantic levels, AMPFlow demonstrates the highest JS-6 score and the lowest ESM-2 pppl. These highlight ProtFlow’s ability to capture the characteristics of real AMPs, in terms of physicochemical properties, sequence composition and semantic meanings.

**Table 2:**
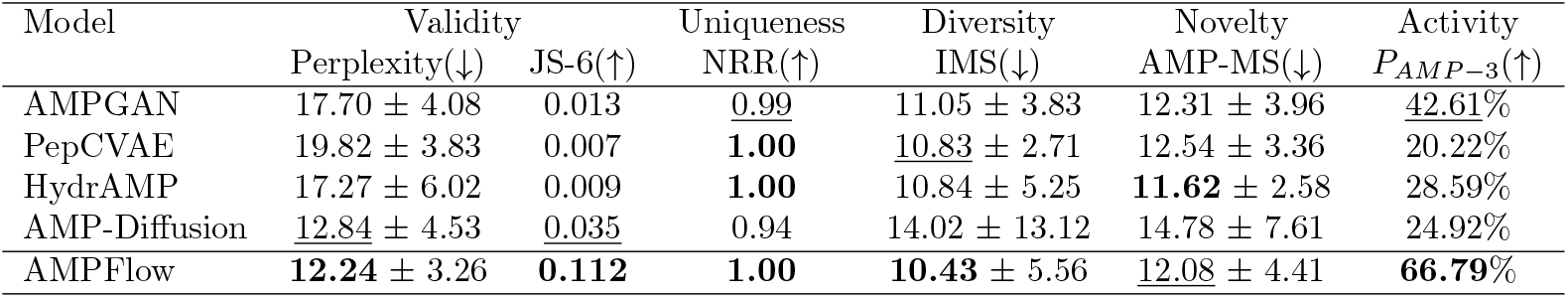
Comparison of computational properties measuring validity, uniqueness, diversity, novelty and activity between ProtFlow and baseline models on the AMP dataset. The **best** and <u>second best</u> scores are marked. *P*_*AMP* −3_ for real AMPs and Uniprot peptides are 68.74% and 0.72% respectively.

**Figure 3.**
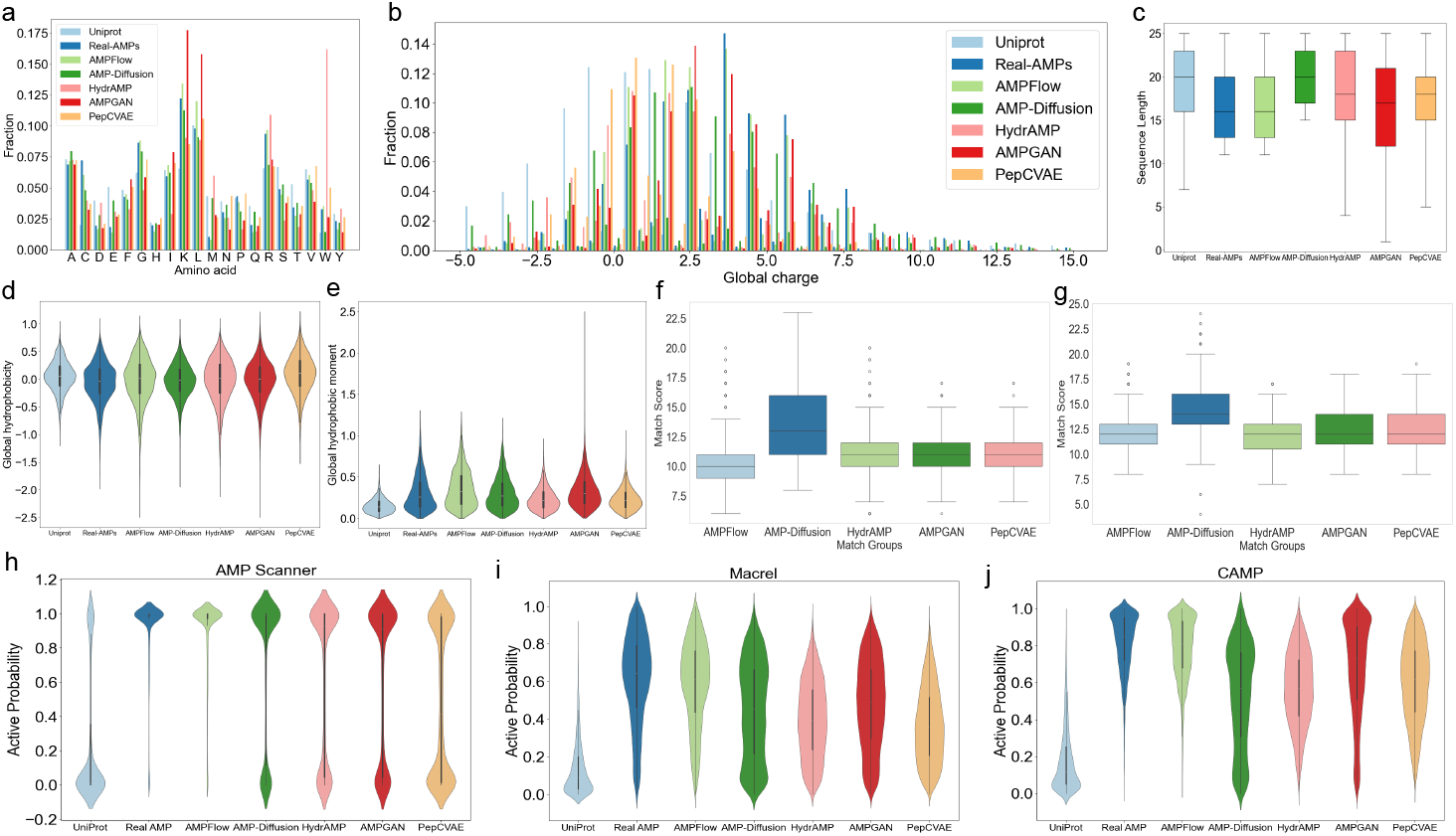
Comparison of physicochemical properties on AMP datasets. (a) Amino acid frequency distribution; (b) Global charge distribution; (c) Sequence length distribution; (d)Eisenberg hydrophobicity distribution; (e) Eisenberg hydrophobic moment distribution; (f)Internal match score distribution within designed peptides; (g)AMP match score distribution between designed peptides and real AMPs; (h)-(j) Activity probability distribution predicted by three AMP predictors.

We further analyzed of the uniqueness, diversity, and novelty of the peptides generated. For uniqueness, we calculated the proportion of non-redundant sequences in the generated data (Non-redundant Ratio, NRR). Since the evaluation of diversity and novelty is not straightforward, we introduced two metrics: Internal Match Score (IMS) and AMP Match Score (AMP-MS), which are defined as the match score between each sequence generated by a method and all the other sequences in the same generated set or real AMP sequences respectively, and retaining the highest score. A lower value indicates fewer overlaps between the source sequence and targeted dataset, reflecting better diversity and novelty.

Since the match score is directly related to the length distribution of sequences, we evaluated sequences with lengths between 10 and 25 across all models for fairness. As shown in Figure 3f-g and Table 2, most methods, including ProtFlow, generate no duplicate sequences, while our method demonstrates the lowest IMS and the second lowest AMP-MS. This observation reflects a trade-off between novelty and similarity. By learning in the semantic space of ESM-2 rather than directly learning from the sequences themselves, our method achieves high similarity to natural AMPs while maintaining remarkable novelty. These findings demonstrate that our method not only generates peptides with high similarity, validity, and reliability, but also ensures their uniqueness, diversity, and novelty.

To evaluate the antimicrobial activity of the generated peptides, we utilized three robust public AMP classifiers: AMP-Scanner v2 [50], Macrel [39], and CAMPR4 [14], to determine antimicrobial activities of designed peptides. Following the recommendation of AMP Scanner, we analyzed only peptides longer than 10 to ensure the accuracy of the predictions. Generally, a peptide sequence is considered an active AMP if its score assigned by the predictor exceeds 0.5. Peptides generated by ProtFlow illustrates high similarity of antimicrobial activity to real AMPs across all three classifiers (Figure 3h-j), and noticeably highest percentage the proportion of peptides identified as AMPs by all three classifiers (Table 2, *P*_*AMP* −3_). This analysis highlights that ProtFlow effectively learns the antimicrobial composition of AMPs from the perspective of protein semantics, driven by the strong data distribution fitting capability of FM and the integration of semantic information.

### 3.5 AMPFlow covers high MIC distribution for multiple bacteria

To determine whether AMP designing models also face the distribution centralization issue, we visualized the generated peptides of each model alongside real AMPs using the method in Section 3.2. Consistent with the results observed in the general peptide scenario, we found that only AMPFlow was able to adequately cover the full distribution of real AMPs. AMP-Diffusion covered a relatively wide distribution range, but still left significant areas uncovered, while the remaining three models were restricted to very limited regions (Figure 4a-e).

**Figure 4.**
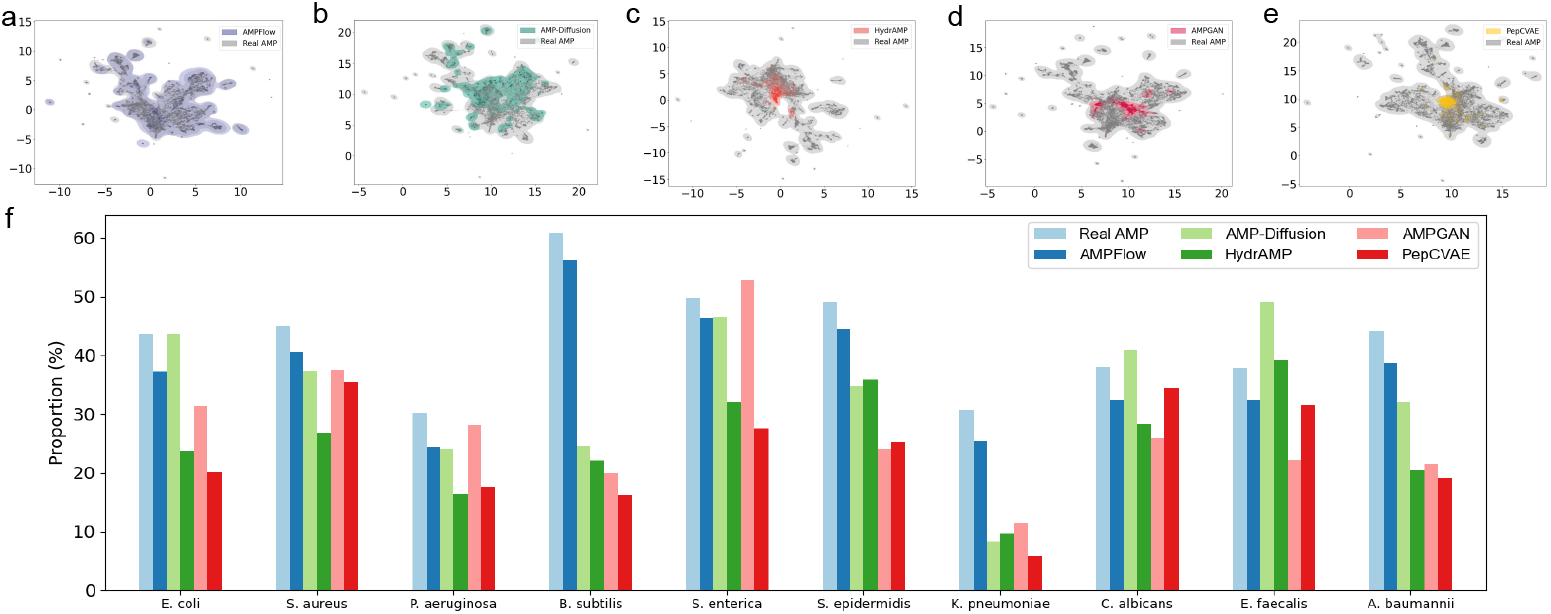
Comparison of protein semantical coverage and multi-bacteria minimum inhibitory concentration (MIC) distributions. (a)-(e) Real AMP distributional coverage in protein semantical embedding space. (f) Proportion of high MIC peptides identified by each bacterium-specific MIC predictors.

To analyze the real-world implications of this issue in computationally assisted AMP design, a straightforward approach is to examine the antimicrobial activity of generated AMPs against different bacterial strains. Because our analysis is based on the protein semantic space, the specificity of AMPs against different bacteria, as a form of protein semantic feature, is expected to exhibit differentiation within this space. The inability of a model to adequately cover the AMP distribution, resulting in the omission of special or rare proteins, may manifest as the model’s inability to generate peptides with strong activity against certain bacterial strains, or as a lower proportion of such peptides in its output. To test this hypothesis, we trained regression models for the top 10 most abundant microorganisms in the MIC dataset. We then used these regressors to predict the logMIC values of peptides from the real AMP dataset and the peptides generated by each model for each bacterial species. Following the settings of [46], we classified peptides with logMIC ≤ 1.5 (MIC ≈ 32 *µ*g/mL) as highly active (positive) and those with logMIC *>* 1.5 as low active (negative). Finally, we calculated the proportion of active peptides against each bacterial species for each model. As illustrated in Figure 4f, the proportion of ProtFlow generated peptides active against these ten microorganisms closely align with the pattern of real AMP, achieving on average 87.46% of the proportion. For *B. subtilis, K. pneumoniae* and *A. baumannii*, we observed that the proportion of active peptides generated by the baseline methods is significantly lower than that found in real AMPs. These observations are consistent with our analysis and hypotheses regarding distribution coverage. These results indicates that our method effectively simulates and covers the AMP space, generating a relatively abundant and reliable pool of candidates.

## 4 Discussion and Conclusion

In this study, we present ProtFlow, a flow matching-based framework for *de novo* protein sequence design. Our method efficiently generate realistic, diverse, and novel peptides and AMPs with broad spectrum activity, especially against previously under-represented pathogens.

One of ProtFlow’s key innovations lies in its ability to leverage semantic information from large protein language models alongside flow matching, allowing for more comprehensive coverage of the training distribution. This stands in contrast to many existing methods, which predominantly generate sequences from high-probability regions, ignoring rare or under-represented clusters of the distribution. Consequently, ProtFlow is able to produce a greater diversity of functional sequences, addressing an unmet need in antimicrobial design.

ProtFlow also utilizes a redesigned, compact, and smooth semantic space to enable faster and more robust training, while retaining biologically realistic sequences. Our algorithm efficiently generates new peptides and AMPs in just a few sampling steps, outperforming state-of-the-art methods both in sampling speed and coverage of the training distribution. This combination lets us identify a larger pool of potent antimicrobial candidates with desirable activity profiles, a crucial advantage for developing broad-spectrum antibiotics against growing resistance. While we cannot directly evaluate performance against future AMR strains, the ability of our model to generate effective candidates for under-represented pathogens suggests enhanced readiness for addressing some of the new threats as they arise.

ProtFlow can serve as a powerful foundation model under the new FM paradigm. However, the model currently operates in an unconditional setting; it can be naturally extended to conditional generation by incorporating auxiliary conditioning variables, such as functional labels or physicochemical constraints, into the flow dynamics, providing a flexible pathway toward controllable protein design.

While this work focuses on antimicrobial peptides, ProtFlow is broadly applicable to other protein design tasks, including signal peptides, antibodies, and enzyme variants, especially in scenarios with limited or unevenly distributed training data. At the same time, several extensions remain to be explored. Future work will focus on scaling the framework to longer protein sequences, as well as more complex settings such as multi-domain and multi-chain proteins. Furthermore, our approach can be naturally integrated with structural information or other modalities, opening up additional possibilities for multiscale, data-informed protein design. Extending ProtFlow in these directions represents an important step toward general-purpose protein sequence generation across diverse biological contexts.

## Acknowledgements

This research was partially supported by “Pioneer” and “Leading Goose” R&D Program of Zhejiang under Grant No. 2024C03048, National Natural Science Foundation of China under Grant No. T2541004, State Key Laboratory of Transvascular Implantation Devices under Grant No. SKLTID2024003 and TIDRI under grant No. KY052025003.

## Source Code Availability

All codes are available at https://github.com/HiracharleFranklin/ProtFlow.

