## Supplemental File for "ProtFlow: Flow Matching-based Protein Sequence Design with Comprehensive Protein Semantic Distribution Learning and High-quality Generation"

### Appendix

#### A Model Architecture

For the decoder, we employ the same architecture as the ESM-2 LMHead, which consists of two linear layers, an activation layer, and a layer norm.

For the compression and decompression modules, we made modifications based on the settings of Hourglass Transformer [18]. For the main experiment with the embedding of ESM-2 35M, we utilize the hidden size of 480, the depth of 4, 8 attention heads, 64 dimensional heads and no hidden dropout rate. The pooling layer uses mean pooling and the activation layer uses tanh function. The linear layers make projections between hidden size of 480 and the compressed hidden size of 30. The two additional sets of normaliztion and denormaliztion layers are dimensional z-score and min-max normalizations. For ESM-2 8M and ProtT5, we set the hidden size to 320 and 1024, respectively.

The backbone of the flow matching holder module is a 12-layer BERT model with 16 attention heads. We use the GELU activation function, with an intermediate layer size of 3072 and an attention dropout rate of 0.1. Depending on the pLM variant employed, the hidden size is set to 320 when using ESM-2 8M, 480 when using ESM-2 35M and 1024 when using ProtT5, with a hidden dropout rate of 0.1. To match the compressed hidden size of the compressor and decompressor, we add two additional linear layers to make projections between hidden size of 480 and the compressed hidden size of 30. The noised protein sequence embeddings are added with positional encodings before being fed into the flow matching holder module. Timesteps are projected to match the size of the protein sequence embeddings using a sinusoidal embedding block, and these are added to the input of each Transformer block after a linear projection. We maintain the advanced long skip connections from [21], which involve adding the linear projections of earlier block inputs to those of later blocks.

#### B Experimental Settings

For fine-tuning the ESM-2 decoder, the model is trained for 1 epoch with 1000 steps, and validation is performed every 50 steps. The learning rate is set to 0.00005. We use a batch size of 512 and the AdamW optimizer, with beta values of (0.9, 0.98) and a weight decay of 0.001.

For training of the compressor and decompressor, all experiments converge within 30 epoches. We perform validation using reconstrction accuracy on the validation set and save checkpoints every 5000 iterations. A cosine learning rate scheduler with two cycle limit is employed, featuring a minimum learning rate of 8e-5 and 10000 linear warmup steps. The batch size is set to 16 for both training and validation. The AdamW optimizer is configured with beta values of (0.9, 0.999).

For training of the flow matching holder, all experiments converge within 1,000,000 iterations. We perform validation using FPD on the validation set and save checkpoints every 10,000 iterations. A cosine learning rate scheduler with a single cycle limit is employed, featuring a minimum learning rate of 0.0002 and 5000 linear warmup steps. The batch size is set to 64. The AdamW optimizer is configured with beta values of (0.9, 0.98), the weight decay of 0.01, the epsilon of 0.000001, and the gradient clipping norm of 1. We trained on one NVIDIA GeForce RTX 3090 GPU. Training models on the general peptide and AMP task for 1,000,000 training steps takes around 42 hours.

For inference, we test the Euler ODE solver with $N \in \{1, 5, 10, 25\}$ steps for reflow and the Dormand Prince 45 ODE solver with $N \in \{1, 5, 10, 25, 50, 75, 100\}$ steps for the other tasks. The optimal solver and sampling steps are different for different tasks. Normally the Dormand Prince 45 ODE solver with a sampling step of 25 can give relatively good results with robustness. For reflow task, the model can support 1-step generation.

For training of the MIC regressors, we use MSE loss and a constant learning rate 0.0001. The batch size is set to 128. We utilize AdamW optimizer with beta values of (0.9, 0.999), the weight decay of 0.01, and the epsilon of 0.000001. Each regressor is trained for 200 epoches.

#### C Compression Ratio

To address the issue of unnecessarily large dimensions in the encoding space of pLM and make the information in this space more focused and compact, we designed a dimensional compressor-decompressor pair positioned between the encoder and decoder. This mechanism compresses the original continuous embedding $h \in \mathbb{R}^{L \times D}$ to $h_c \in \mathbb{R}^{L \times D/c}$ with a compression ratio $c$ and subsequently reconstructs it back to $h' \in \mathbb{R}^{L \times D}$. To determine the optimal value of the compression ratio $c$, we evaluate the effects of a range of compression ratios $[1, 2, 4, 8, 16, 32, 60, 120, 160]$ by assessing the reconstruction accuracy and FPD.

The results, summarized in Table S1, demonstrate that when the dimensional compression ratio is less than 32, the model can accurately reconstruct the sequence without significant differences, while the reconstruction accuracy begins to decline when the compression ratio is greater than 32, until the sequence can no longer be effectively reconstructed at the ratio of 160. As the embedding dimensionality is progressively reduced up to a ratio of 16, the FPD values improve in both cases, indicating an enhanced capability for distribution learning. Beyond this threshold, additional compression may introduce potential trade-offs between efficiency and representation fidelity, causing substantial performance degradation. In general, the model shows strong robustness to the ratios between 1 to 60.

Table S1: Reconstruction accuracy and FPD of each Compression Ratio (c) on the general peptide and long-chain protein tasks.

| Ratio (c) | Acc(%)↑ | FPD↓ |
|---|---|---|
| 1 | 99.93 | 0.48 |
| 2 | 99.96 | 0.44 |
| 4 | 99.97 | 0.42 |
| 8 | 99.96 | 0.41 |
| 16 | 99.96 | 0.36 |
| 32 | 99.98 | 0.43 |
| 60 | 98.87 | 0.46 |
| 120 | 92.79 | 0.60 |
| 160 | 26.53 | - |

### D Ablation Study

ProtFlow, as a method based on the latent encoding space of protein language models (pLMs), places particular emphasis on the impact of the quality of the encoding space on model performance. Moreover, the performance of the encoder and decoder may directly determine the upper limit of latent space-based models. Therefore, we first evaluated three different protein language model encoders, ESM-2 8M, ESM-2 35M and ProtT5-XL [8], on the general peptide dataset. This settings includes two different series of pLMs with different parameter acquisitions. Besides, the introduction of ProtT5 which can eliminate the possible advantages caused by ESM-2 in terms of ESM-based metrics ESM-2 pseudoperplexity and ESM-IF scperplexity. Specifically, we calculated the reconstruction FPD for these pLMs: ESM-2 8M scored 0.03592, while ESM-2 35M scored 0.01487 and ProtT5 scored 0.00405, indicating that ESM-2 35M and ProtT5 demonstrate increasingly superior protein sequence reconstruction capability. As shown in Table S2, the two ESM-2 encoders performed comparably on metrics reflecting the quality of sequence and structure generation (pLDDT, ESM pppl, ESM-IF scPerplexity, and TM-Score), with minor differences in their scores, while ProtT5 leads slightly in all these metrics. This suggests that these encoders are sufficiently capable of enabling ProtFlow to learn high-quality protein sequences, but encoders with better representation and reconstruction abilities can generate better results. Notably, experiments based on ESM-2 35M consistently outperformed ESM-2 8M across FPD, MMD, and OT metrics, which reflect the ability to learn data distribution, and these based on ProtT5 outperformed ESM-2 35M. This demonstrates that the quality of the protein language model's encoding space has a significant impact on ProtFlow's ability to learn the distribution of protein space.

We conducted a redesign of the ESM-2 35M latent encoding space, incorporating compression, smoothing, and normalization operations. ProtFlow, based on the redesigned protein language model latent space, demonstrated consistent performance improvements across metrics reflecting peptide design reliability, foldability, naturalness, and three distribution similarity measures. This highlights that our redesign module significantly enhances the quality of the protein language model's latent space. It not only increases the information density of the space and aligns the distribution closer to a standard Gaussian distribution but also regularizes outlier subgroups. Consequently, this optimization improves ProtFlow's performance in both protein sequence design based on semantic understanding and the fitting of protein semantic space distributions.

Table S2: Ablation Study with Different encoders/decoders and the reflow technique.

| Encoder | Decoder | Reflow | pLDDT (↑) | ESM-2 pppl (↓) | scPerplexity (↓) | TM-score (↑) | FPD (↓) | MMD (↓) | OT (↓) | NFE (↓) |
|---|---|---|---|---|---|---|---|---|---|---|
| ESM-2 8M | - | No | 71.14 | 12.89 | 13.05 | 0.56 | 0.53 | 0.010 | 2.07 | 25 |
| ESM-2 35M | - | No | 70.54 | 12.96 | 12.83 | 0.56 | 0.46 | 0.009 | 1.99 | 25 |
| ProtT5 | - | No | 71.54 | 12.76 | 12.74 | 0.57 | 0.44 | 0.007 | 1.90 | 25 |
| redesigned ESM-2 35M | linear-based | No | 70.92 | 12.92 | 12.99 | 0.54 | 0.57 | 0.009 | 2.15 | 25 |
| redesigned ESM-2 35M | Transformer-based | No | 71.93 | 12.29 | 12.14 | 0.59 | 0.36 | 0.004 | 1.86 | 25 |
| redesigned ESM-2 35M | Transformer-based | Yes | 72.86 | 12.16 | 11.79 | 0.60 | 0.45 | 0.006 | 1.97 | 1 |

To estimate the impact of the decoding block architecture on the model performance, we ablated the Transformer-based decompression network with a linear layer. This modification leads to inaccurate sequence reconstruction with a 0.9342 accuracy, and further results in a comprehensive reduction of all metrics. This set of experiments proved the important role of a reasonable and precise reconstruction architecture in latent space learning.

One of the most significant advantages of FM is its fast generation speed. Combining the observations in Table S1 and Table S2, unlike diffusion models, which require hundreds or even thousands of steps, Dirichlet FM achieves comparable results with only 50 steps, making it twice as efficient as DiMA. Leveraging the nearly linear probability path of 1-RF, ProtFlow achieves excellent results with just 25 steps, achieving a fourfold speed improvement over DiMA. Reflow (2-RF) further straightens the probability path, enabling even single-step generation using the simplest Euler ODE solver. Although fine-tuning based on imprecise 1-RF probability path sampling may slightly compromise distributional performance, reflow still outperforms all other methods. Notably, the reflow operation further improved the performance of ProtFlow across pLDDT, ESM pppl, ESM-IF scPerplexity, and TM-Score. We hypothesize that this improvement is likely due to reflow's ability to straighten probabilistic paths, thereby reducing intersections between different probability paths. This reduction in path overlap minimizes data ambiguity, which in turn facilitates learning the composition and semantics of individual protein sequences more effectively.

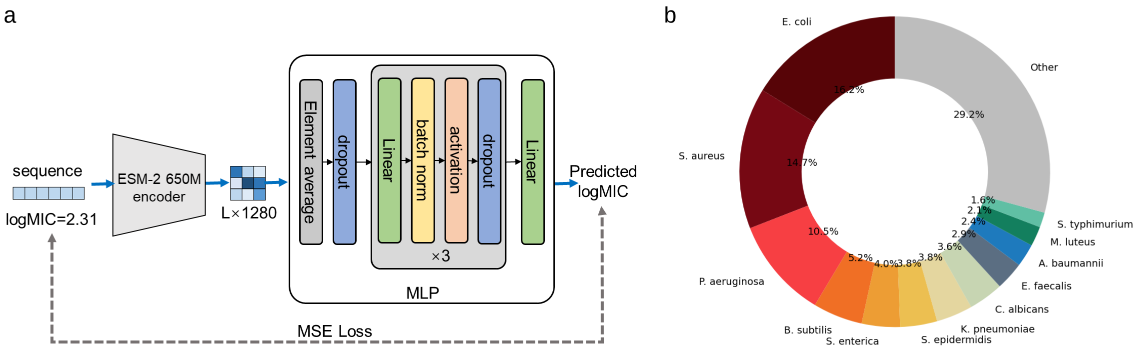


Figure S1: Architecture of the MIC regressor and bacteria proportion in the MIC dataset. (a) Architecture of the MIC regressor. (b) Proportions of leading bacteria in the MIC dataset.

### E MIC Regressors

As shown in Figure S1a, the MIC predictor we constructed consists of a protein language model encoder and a modified multilayer perceptron (MLP). We use the ESM-2 650M encoder to transform input sequences into embeddings of size $L \times 1280$, incorporating rich protein semantic information through the protein language model during this process. The MLP is composed of three layers, each consisting of a linear layer, a batch normalization layer, a ReLU activation layer, and a dropout layer. The first layer reduces the embedding dimension from 1280 to 1024, and each subsequent layer halves the input dimension. Finally, an output layer predicts the logMIC with a single output dimension. Additionally, we include an element-wise average layer with dropout before the MLP. This layer compresses the $L \times 1280$ embeddings into $1 \times 1280$ to integrate global sequence information, with the dropout rate set to 0.1.

The fraction of AMPs targeting different bacteria in our collected MIC dataset is summarized, as shown in Figure S1b. Among these, only *E. coli*, *S. aureus*, and *P. aeruginosa* are considered common bacteria, with each corresponding to over 5,000 AMPs, accounting for more than 10% of the total dataset. Beyond these three, the number of AMPs targeting other bacteria drops sharply. For instance, *B. subtilis*, ranked fourth, has only 2,666 AMPs, representing 5.2% of the dataset. When the number of AMPs is too small, the predictions from the trained regression model may lack reliability. Therefore, we selected the top 10 bacterial strains(*E. coli*, *S. aureus*, *P. aeruginosa*, *B. subtilis*, *S. enterica*, *S. epidermidis*, *K. pneumoniae*, *C. albicans*, *E. faecalis*, *A. baumannii*) to form an MIC subset, which includes three common bacteria and seven rare bacteria. This subset comprises 4 Gram-positive bacteria: *S. aureus* (7597 entries), *B. subtilis* (2666 entries), *S. epidermidis* (1987 entries), *E. faecalis* (1523 entries); 5 Gram-negative bacteria: *E. coli* (8372 entries), *P. aeruginosa* (5417 entries), *S. enterica* (2054 entries), *K. pneumoniae* (1957 entries), *A. baumannii* (1233 entries) and 1 fungus, *C. albicans* (1871 entries), providing a balanced representation of both abundant and scarce bacterial types as well as various bacterial types with specific characteristics.

For training of the MIC regressors, we use MSE loss and a constant learning rate 0.0001. The batch size is set to 128. We utilize AdamW optimizer with beta values of (0.9, 0.999), the weight decay of 0.01, and the epsilon of 0.000001. Each regressor is trained for 200 epoches. We evaluated the predictive performance of our regression model using Root Mean Squared Error (RMSE) and Coefficient of Determination ($R^2$). RMSE directly reflects the magnitude of the prediction error between the predicted and true values, while $R^2$ measures the correlation between the predicted and true values. A smaller RMSE and a larger $R^2$ indicate more accurate MIC predictions. First, we compared the performance of our model with an RNN-based model and a model where the MLP was replaced by LSTM, using the most common dataset for *E. coli.* Our model achieved an RMSE of 0.2772 and an $R^2$ of 0.8805, outperforming the RNN model (RMSE: 0.5548, $R^2$: 0.5126) and the LSTM-based model (RMSE: 0.4137, $R^2$: 0.7344). Subsequently, we trained an AMP MIC predictor for each microbial species. As shown in Table S3, the RMSE values for all species were below 0.3, and the $R^2$ values exceeded 0.85, demonstrating consistent predictive accuracy across different microbial species.

Table S3: Performances of MIC predictors

| Bacteria | RMSE↓ | $R^2$↑ |
|---|---|---|
| *E. coli* | 0.2772 | 0.8805 |
| *S. aureus* | 0.2683 | 0.8759 |
| *P. aeruginosa* | 0.2554 | 0.8815 |
| *B. subtilis* | 0.2335 | 0.9168 |
| *S. enterica* | 0.2346 | 0.9188 |
| *S. epidermidis* | 0.2235 | 0.9083 |
| *K. pneumoniae* | 0.2254 | 0.9203 |
| *C. albicans* | 0.2627 | 0.8829 |
| *E. faecalis* | 0.2401 | 0.9096 |
| *A. baumannii* | 0.2115 | 0.9243 |

### F Baseline Models

#### F.1 Baselines for General Peptide Design

We use general peptides, which are more representative and broadly applicable, as a case study to discuss the algorithmic innovations of our method. Our comparison comprehensively covers the major paradigms of current generative models, including GANs, autoregressive (AR) models, discrete diffusion models, latent diffusion models, and discrete flow matching, represented by ProteinGAN, ProtGPT2, EvoDiff, DiMA, and Dirichlet Flow Matching, respectively. For a fair comparison, most models are evaluated using versions with parameter scales comparable to ours. At the same time, considering the widespread adoption of large-scale models in current research, we also include ProtGPT2 with a substantially larger number of parameters. Detailed descriptions of each model are provided below.

**ProteinGAN** represents a pioneering application of generative models in *de novo* protein sequence design. It is a variant of the generative adversarial network (GAN), where both the discriminator and generator are based on ResNet-based convolutional neural networks (CNNs), further enhanced with a self-attention layer. While ProteinGAN was originally developed for Multiple Sequence Alignment

(MSA) tasks, which focus on specific protein families, it has also demonstrated competitive performance in our general peptide design task.

**ProtGPT2** is a groundbreaking autoregressive language model designed for *de novo* protein sequence generation, leveraging the powerful GPT-2 architecture to model the protein sequences. Specifically, ProtGPT2 leverages the GPT-2 LMHead model architecture containing 36 layers with a model dimensionality of 1280, totally 738 million parameters. We apply full-parameter finetuning to the standard ProtGPT2 on our dataset.

**EvoDiff-OADM** is the first foundation diffusion model for protein design trained on evolutionary-scale protein sequence data. EvoDiff grounds in two famous discrete diffusion frameworks: D3PM and OADM, while the OADM-based model is reported to perform better in the original paper. EvoDiff-OADM operates in an order-agnostic autoregressive manner, gradually converting single amino acids to or from the mask token during the forward or reverse processes. The model is based on the ByteNet CNN architecture, and we utilize a configuration with 38 million parameters. Furthermore, we evaluate its performance after replacing the backbone with the Transformer-based ESM-2 35M architecture.

**DiMA** is one of the earliest diffusion model operating on continuous space by embedding discrete protein sequences with the protein language model ESM-2 8M and surpasses lots of traditional mainstream generative models. It is built on a 12-layer transformer model and introduces multiple novel techniques to improve the performance, including long skip connections, self-conditioning and tanh noise schedule.

**Dirichlet Flow Matching** is a pioneering approach in discrete flow matching, framing generative modeling as a transportation problem on the probability simplex and using the Dirichlet distribution as the probability distribution for the generative path. It has demonstrated strong performance in DNA sequence design tasks. We adapted Dirichlet Flow Matching from the DNA sequence design into the amino acid sequence generation of protein, retaining the original CNN architecture and model hyperparameter settings.

#### F.2 Baselines for AMP design

We use antimicrobial peptides, one of the most extensively studied classes of therapeutic proteins, as a representative case study to demonstrate the application of our algorithm to functional peptide design. Given the specialized nature of designing proteins for specific functions, we include several mainstream *de novo* antimicrobial peptide design models for comparison, including HydrAMP, PepCVAE, AMPGAN, and AMP-Diffusion. To ensure a fair comparison, most baseline models are evaluated using versions with parameter scales comparable to our model. Detailed descriptions of each model are provided below.

**HydrAMP** is based on a conditional variational autoencoder (cVAE) with an autoencoder and a decoder, and incorporates a pre-trained classifier. It captures the antimicrobial properties of peptides by learning their low-dimensional, continuous representations and decouples these properties from the antimicrobial conditions. The model is designed to generate peptide sequences that meet specific antimicrobial activity criteria. HydrAMP can perform both unconstrained generation and generate antimicrobial analogs based on provided prototype peptides.

**PepCVAE** is a semi-supervised generative model also based on a cVAE and incorporates a pre-trained Antimicrobial Peptide (AMP) classifier. The model learns a rich latent space representation by leveraging a large dataset of unlabeled peptide sequences along with a smaller set of sequences labeled as either antimicrobial or non-antimicrobial. By decoupling the antimicrobial properties from the latent space, PepCVAE is able to generate peptide sequences that meet specific antimicrobial objectives.

**AMPGAN** is based on a Bidirectional Conditional Generative Adversarial Network (BiCGAN). In addition to the standard generator and discriminator components found in typical GANs, it introduces an encoder that maps real data into the generator's latent space. This enables iterative peptide sequence generation, optimization, interpolation, and incremental modifications. The generation process in AMPGAN is controlled by conditioning variables, allowing the discriminator to learn the relationship between relevant antimicrobial features and the generated peptide sequences.

**AMP-Diffusion** is built on the continuous diffusion model framework, where peptide sequences are mapped into a latent space using the protein language model ESM-2 for the diffusion process. The model architecture incorporates pre-trained ESM-2 8M attention blocks for the denoising process and a multilayer perceptron (MLP) as the output layer. The timestep is embedded with a positional encoding and integrated into protein embeddings with a scaling factor and a bias adjustment. AMP-Diffusion reports a leading performance compared to mainstream AMP design methods and is able to generate AMPs with good predicted activities.

The comparative methods included in this study case are primarily based on VAE- and GAN-style generative frameworks, which currently dominate *de novo* antimicrobial peptide design. This selection reflects common practice in the field, where such models have been most extensively explored and validated for antimicrobial peptide generation. Accordingly, these methods serve as representative and well-established baselines for evaluating new approaches.

### G Metrics Calculation

**ESM-2 pseudoperplexity (ESM-2 pppl)** measures how well a given protein sequence aligns with the patterns the assessing model has learned from the training data of ESM-2, which is a large-scale standard protein database UniRef50 [27]. We calculate ESM-2 pppl with ESM-2 35M [15] by masking each amino acid of the protein sequence and predicting it considering all the other amino acids in the sequence, and calculating the value with the equation

$$\text{ESM-2 pseudoperplexity} = \exp\left(-\frac{1}{|x|}\sum_{i=1}^{|x|}\log p(x_i \mid x_{j\neq i}, \theta_{\text{ESM-2}})\right)$$

where $|x|$ means the length of sequence x and $x_{j\neq i}$ means sequence x without the $i^th$ amino acids.

**TM-Score** [33] evaluates the similarity between two protein pairs in structure. Compared to pLDDT which also make evaluations on the structure level, TM-Score focuses more on the global level. For each protein sequence, after predicting the structure with OmegaFold [30], we use the FoldSeek [12] easy-search tool to seek for the most similar protein in the AlphaFold SwissProt [29] database with the TM-Score function

$$\text{TM-score} = \frac{1}{|x_{query}|}\sum_{i=1}^{|x_{query}|}\frac{1}{1+\left(\frac{d_i}{\sigma|x_{target}|}\right)^2}$$

where $|x_{query}|$ means the number of aligned residues between the query and target proteins, $|x_{target}|$ is the length of the target protein, $\sigma$ is a scaling factor, and $d_i$ is the distance between the $i^{th}$ aligned residue pairs.

**Fréchet ProtT5 Distance (FPD)** is a variant of Fréchet distance (FD) [2], which measures the dissimilarity between two samples drawn from multivariate Gaussian distributions. Given two samples $X_1 \sim \mathcal{N}(\mu_1, \Sigma_1)$ and $X_2 \sim \mathcal{N}(\mu_2, \Sigma_2)$, the FID can be calculated as

$$\text{Fréchet Distance} = \|\mu_1 - \mu_2\|^2 + \text{tr}(\Sigma_1 + \Sigma_2 - 2\sqrt{\Sigma_1\Sigma_2})$$

We calculate the Fréchet distance of protein sequence embeddings using protein language model ProtT5, namely, Fréchet ProtT5 Distance (FPD) [1].

**Maximum mean discrepancy (MMD)** [23] is a kernel-based statistical test to determine whether two samples $X = \{x_1, x_2, \ldots, x_n\}$ and $Y = \{y_1, y_2, \ldots, y_n\}$belong to different distributions. Suppose the kernel is k, then MMD can be calculated as

$$\text{MMD} = \frac{1}{n^2}\sum_{i=1}^{n}\sum_{j=1}^{n}(k(x_i, x_j) + k(y_i, y_j) - 2k(x_i, y_j))$$

We calculate MMD on the ProtT5 embeddings and use the radial basis function kernel.

**1-Wasserstein optimal transport (OT)** [24] evaluates the similarity between two batches of sequences. We use pairwise Levenshtein distances [22] as transportation costs, utilize the Earth Mover Distance (EMD) solver [3] with a uniform distribution of the samples to determine optimal sequence pairs, and take the average of the distances between optimal pairs.

**Jaccard Similarity Coefficient** is a widely used measurement to evaluate the similarity of two sets. We use 6-mers Jaccard similarity coefficient (JS-6) to compare the similarity of generated peptide sets to the training data. We split the training dataset and the generated peptide set with 6-mers, which represents all the sub-sequences with length 6 in the dataset to be processed. Then the Jaccard similarity can be calculated with the equation

$$\text{Jaccard}(A, B) = \frac{|A \cap B|}{|A \cup B|}$$

where A represents the 6-mers set of training dataset and B is the 6-mers set of generated batch set.

**Match Score** In this study, we includes Internal Match Score (IMS) and AMP Match Score (AMP-MS). Both metrics are based on the *pairwise2.align.globalxx* tool of *biopython* [5] for the calculation of the match score, representing the number of matching elements between a pair of sequences. Higher match scores indicate greater overlap in composition. For IMS, we calculate the match score between each sequence generated by a method and all the other sequences in the same generated set, retaining the highest score. This value measures the degree of repetition within the dataset itself, where a lower score reflects greater diversity in the generated peptide set to some extent. For AMP-MS, we calculate the match score between each sequence generated by a method and the real AMP sequences, retaining the highest score. A lower value indicates that the generated peptides do not simply replicate existing sequences or sequence fragments from real AMPs, reflecting its novelty. It is important to note that AMP-MS reflects the degree of global overlaps while the previously used JS-6 metric focuses on local functional regions. Therefore, the two metrics may not necessarily show consistency.

### H Physicochemical Analysis for Protein Space

To reveal the inherent differences in protein distribution and thereby provide possible factors that traditional generative models cannot fit into the overall protein space, we comprehensively analyzed the physicochemical property differences between the core and peripheral regions of natural proteins.

For generic peptides, sequences located in peripheral regions exhibit physicochemical properties that differ substantially from those in the core of the distribution (Figure S2). Peripheral peptides exhibit substantially broader length and molecular weight distributions, encompassing both unusually short and extended sequences, whereas core peptides are concentrated around intermediate lengths. In addition, peripheral peptides display more extreme isoelectric points, including strongly basic or acidic sequences, in contrast to the predominantly near-neutral or mildly basic core population. Hydrophobicity is also markedly elevated in the peripheral region, accompanied by enrichment of hydrophobic residues, while core peptides show more moderate and balanced compositions. Furthermore, peripheral peptides tend to exhibit stronger and more homogeneous secondary-structure preferences, particularly increased $\alpha$-helical propensity, whereas core peptides display greater structural heterogeneity. Together, these observations indicate that the peripheral region corresponds to low-density tails of the peptide distribution characterized by extreme and heterogeneous physicochemical properties rather than random noise.

A distinct yet conceptually consistent pattern is observed for AMPs (Figure S3). While AMP sequences overall conform to canonical characteristics such as short length, high positive charge, and amphipathic composition, a clear separation remains between core and peripheral regions. Peripheral AMPs tend to be slightly longer but display more concentrated length distributions, suggesting increased structural consistency. Notably, these peptides are strongly enriched for high isoelectric points, reflecting elevated net positive charge, a defining feature of membrane-active antimicrobial function. Unlike generic peptides, peripheral AMPs do not simply exhibit increased hydrophobicity; instead, they show a more optimized balance between hydrophobic and charged residues, consistent with canonical amphipathic $\alpha$-helical antimicrobial peptides. Moreover, peripheral AMPs demonstrate higher $\alpha$-helical content and lower instability indices, indicating enhanced structural stability. Their amino acid composition further reflects enrichment of helix-favoring and positively charged residues, alongside depletion of residues associated with structural disruption. Importantly, these properties suggest that peripheral AMPs represent a functionally refined and structurally coherent subset rather than statistical outliers.

Despite functional differences between generic peptides and antimicrobial peptides, their peripheral regions share common features, including extreme physicochemical properties, skewed amino acid compositions, and strong structural regularity. These characteristics place peripheral sequences in low-density regions of the overall distribution, which may cause the potential to be systematically underrepresented for generative models.

### I Related Work

***De novo* protein sequence design** focuses on constructing protein sequences that belong to biologically active groups, such as enzymes [31], antibodies [9, 20], and peptides [4] from scratch. Deep generative models have proven effective in designing high-quality *de novo* protein sequences. Prominent methods include protein language models [14] and discrete diffusion models [1]. By encoding proteins

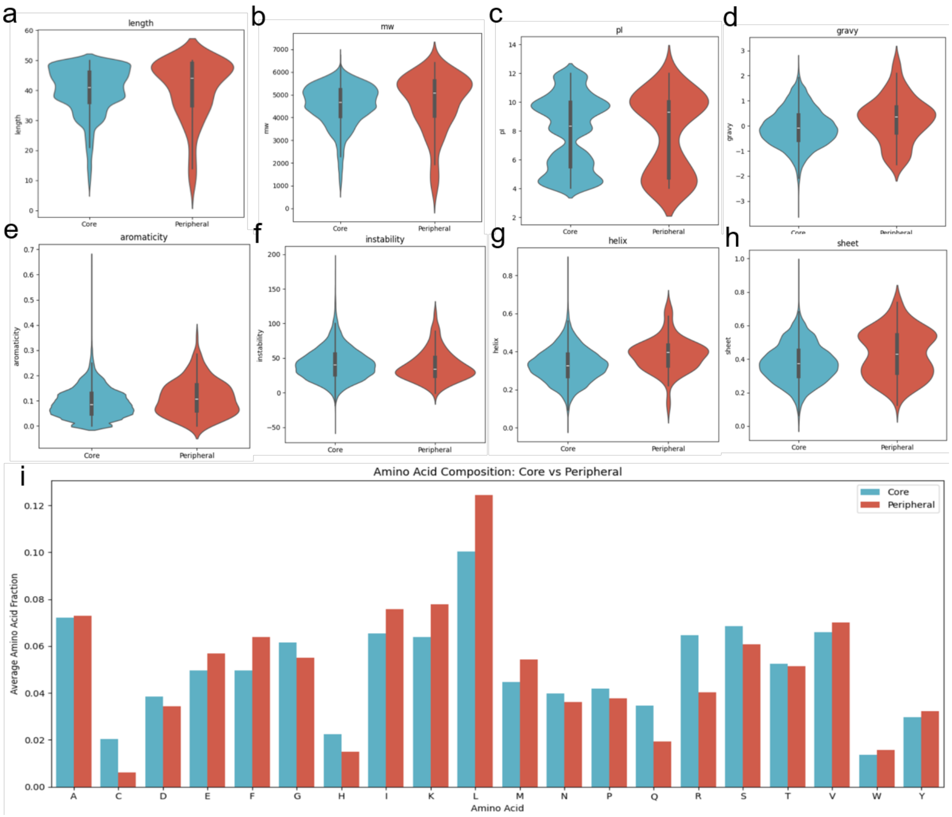


Figure S2: Physicochemical analysis for the core and peripheral region of general peptides. (a)-(h) The length, mw, pI, GRAVY, aromaticity, instability, $\alpha$-helix, $\beta$-sheet distributions; (i) the amino acid proportion distribution.

into a latent space, advanced continuous diffusion models can be effectively applied to this task [21]. Our work leverages this continuous approach to delve into the semantically rich latent protein space.

**Diffusion models** [10] can generate meaningful output by gradually denoising samples from prior noise distributions. To improve efficiency, the score-matching paradigm [26] is introduced. Diffusion models have become a prominent framework in generative modeling, achieving notable success across various domains such as image synthesis [7], text generation [13], and molecular modeling [11]. Despite their successes, diffusion models often encounter challenges like high computational demands, prolonged generation times, and sub-optimal probability paths [25, 19].

**Flow matching models**, proposed as an efficient, simulation-free method to train Continuous Normalizing Flows (CNFs), eliminate the need for explicit knowledge of the marginal vector field [16]. Despite these advancements, conventional solvers still require extensive function evaluations [6]. To address these challenges, several techniques have been developed to improve performance. Rectified Flows (RFs) [17] employ straight-line simulations of probability paths, simplifying training and accelerating sampling by avoiding complex ODE solvers while preventing the collapse of word embeddings. Conditional Flow Matching (CFM)[28] learns the vector field conditioned on individual data points and incorporates Optimal Transport to ensure optimal probability paths, enhancing training efficiency and reducing inference time. While flow-based methods have shown promising results in the design of biological molecules, including protein structures [32], their application to protein sequence design remains largely unexplored. To bridge this gap, we introduce the first FM-based generative model specifically tailored for protein sequence design.

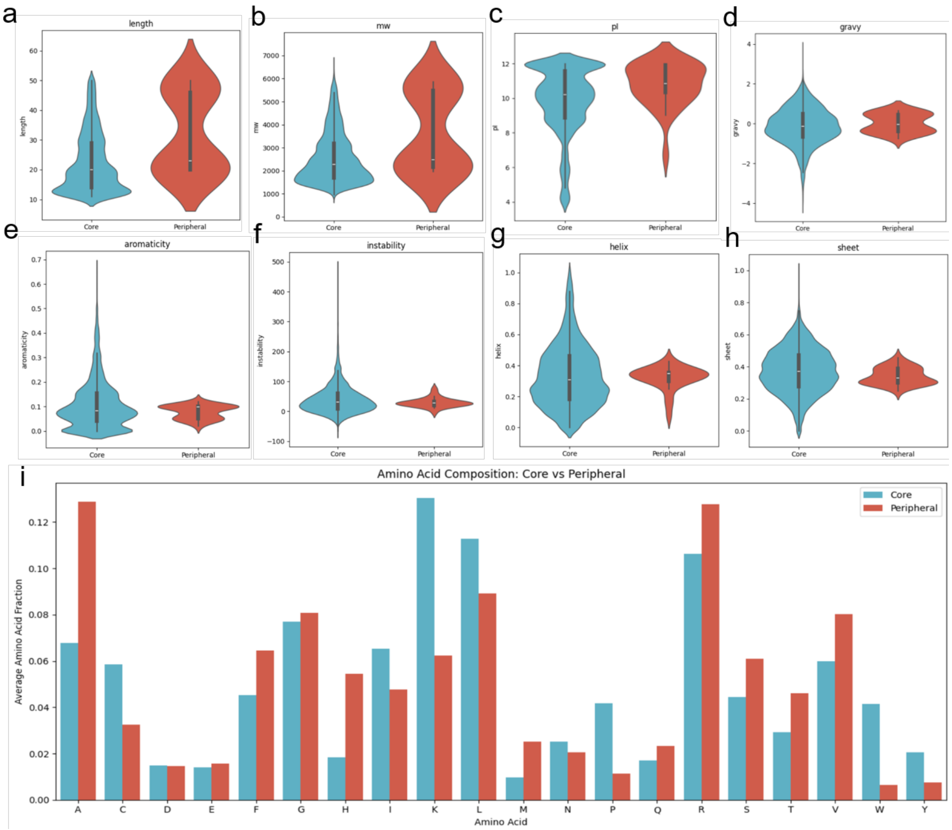


Figure S3: Physicochemical analysis for the core and peripheral region of AMPs. (a)-(h) The length, mw, pI, GRAVY, aromaticity, instability, $\alpha$-helix, $\beta$-sheet distributions; (i) the amino acid proportion distribution.
